# Higher cognitive reserve is associated with better neural efficiency in the cognitive performance of young adults. An event-related potential study

**DOI:** 10.1101/2019.12.30.890830

**Authors:** Gabriela Gutiérrez-Zamora Velasco, Thalía Fernández, Juan Silva-Pereyra, Vicenta Reynoso Alcántara

## Abstract

To examine the effects of cognitive reserve (CR) and working memory (WM) load on the cognitive performance of young adults, we performed two event-related potential (ERP) experiments. The first experiment aims to show how high CR influences young adult performance as a function of two levels of working memory load (high vs. low) during a Sternberg task. For both positive and negative probes, participants with high and low CR showed larger P300 amplitudes to low WM loads than to high WM loads. Both CR groups showed a longer P300 latency to high WM loads than to low WM loads, but this difference was greater for the low CR group than for the high CR group. The high CR group displayed larger P300 amplitudes for every experimental condition compared to the low CR group. The second experiment analyzed grammatical gender agreement in sentence processing when CR and WM load were manipulated. Sentences varied according to the gender agreement of the noun and adjective, where the gender of the adjective either agreed or disagreed with that of the noun (agreement), and with regard to the number of words between the noun and the adjective in the sentence (WM load). Participants with high CR showed greater modulation of left anterior negativity (LAN) and P600a effects as WM increased than that observed in participants with low CR. The findings together suggest that higher levels of cognitive reserve improve neural efficiency, which may result in better working memory performance and sentence processing.

## Introduction

Lifestyle and everyday experiences seem to have a cumulative impact on cognition. An enriched environment during one’s life may play a protective role against cognitive deficits associated with aging [1, 2], which could be a consequence when optimizing the use of resources through the recruitment of neural networks and/or alternative cognitive strategies [2]. This adaptation mechanism, called the cognitive reserve (CR), can be defined as the capacity, flexibility or efficiency of cognitive processes that are associated with the susceptibility to aging or pathology [3, 4]. Although CR cannot be directly measured, it can be assessed by measuring proxy variables such as educational level, intelligence quotient (IQ), occupational complexity and leisure activities that require some intellectual effort [4].

A high CR is associated with good performance in cognitive tasks of working memory, attention and reasoning [5–10]. Its effect on cognition is especially noticeable in older adults and in populations with some brain pathology [11–16]. Thus, CR is considered to increase the efficiency and flexibility of neural networks [17, 18].

Although CR can be observed in the performance of older adults, given the accumulation of experiences from an enriched lifestyle, it can be hypothesized that this phenomenon should be observed even in young adults since CR is present through all stages of development. Thus far, we do not know of any study in which the effects of CR in healthy young adults are evaluated. Studies comparing the performance of older adults with that of young people not only show differences in the processing pattern due to development or age [9, 19], but also in terms of the accumulation of experiences.

Several studies have reported the impact of CR on working memory (WM) [7,9,20–23], since WM represents a measure of fluid intelligence through which CR could act as a protective mechanism of cognitive abilities against cognitive deficits [24, 25]. Based on neuroimaging studies, compensation models have been proposed given the brain activation patterns as an effect of CR during the performance of WM tasks. For example, in the Hemispheric Asymmetry Reduction in Older Adults model (HAROLD model; [26]), it is suggested that neurofunctional changes related to age are associated with a significant reduction of hemispheric lateralization in the prefrontal cortex. Thus, an efficient performance in an older adult that shows bilateral activation can be a reflection of compensation. In contrast, in the Posterior-Anterior Shift in Aging model (PASA model; [27]), the compensation is considered to be the changes in the activation pattern from the posterior to anterior regions associated with aging. In the Scaffolding Theory of Aging and Cognition (STAC, [28]), scaffolds involve the participation of supplementary neuronal circuits that provide the additional computational support required by an older brain to preserve cognitive function in the face of localized or global neurofunctional impairment. In neuroimaging studies, scaffolding is observed as a greater activation or an additional recruitment of the prefrontal and parietal brain regions compared to young adults.

The effects of CR on cognition in younger populations of patients have not been thoroughly studied because specific compensatory mechanisms for these patients have not been proposed from functional studies of electroencephalography or neuroimaging; however, the same mechanisms described in aging could be applied to explain the superiority of patients with greater CR to recover themselves [29, 30].

Studies on Event-related potentials (ERPs) recorded during cognitive tasks allow the gathering of information with great temporal resolution. ERPs are the average of the cerebral electrical activity synchronized to some external or internal events and are classified according to their polarity (i.e., positive or negative deflections in the waves), the time in milliseconds from the stimulus presentation, and their distribution over the scalp. Recently, Gu et al. [6], Speer and Soldan [9], and Sundgren, Wahlin, Maurex and Brismar [31] showed that participants with a higher CR level had a higher percentage of correct answers, shorter response times and a different modulation of the P300 amplitude with respect to subjects with lower CR during a working memory (WM) task with different levels of difficulty. The P300 component is a positive deflection of the ERP with a maximum amplitude at 300 milliseconds and is associated with the updating of working memory [32, 33]. The P300 component has been related to the cognitive demand of a task [33, 34] and the related attentional processes [35]. For instance, Speer and Soldan [9] found that subjects performing a verbal working memory task (the Sternberg task) showed decreasing P300 amplitudes and longer latencies with increasing WM load, but this effect was greater in subjects who had low CR (in both young and older individuals) than subjects with a high CR. This result was interpreted as a greater CR being associated with greater neuronal efficiency in terms of lower neural activity and higher processing speed as the demand for the task increased. Gu et al. [6] reported similar results in healthy adults; they found a negative correlation between changes in the amplitude of the P300 component and the level of CR and suggested that higher CR reduces neural inefficiency. Modulations of the amplitude of P300 have also been associated with a physically active lifestyle [36], educational level [11], and intelligence [37, 38].

To our knowledge, there are few studies on the impact of CR on language, especially on the processing of sentences, perhaps because certain types of processes that are involved, unlike working memory, are associated with crystallized intelligence [39]. Nevertheless, only one behavioral study [40] has examined how print exposure (i.e., habitual investment in reading and literacy activities) affects sentence processing and memory in older adults. They showed that life-long habits of literacy increase the efficiency of component reading processes, buffer the effects of working memory decline on comprehension and contribute to maintaining skilled reading.

Compared with older adults, young people are more cognitively and neurally efficient, but older adults have been shown to be able to compensate, perhaps as mechanisms resulting from CR, although there is no evidence to support this idea. A recent ERP study that analyzed the sentence processing of younger adults with respect to older adults at two levels of WM load by comparing brain electrical activity associated with grammatical gender agreement processing [41] showed a sequence-specific processing pattern in older adults that could reflect the process of compensation. Older adults displayed modulations of the amplitude of the ERPs that denoted an initial failure in the identification of the grammatical violation (i.e., left anterior negativity (LAN), morphosyntactic analysis; [42, 43]) but with a compensation of this process in later stages of the analysis of the sentence (i.e., P600a, integration of arguments and P600b, mapping of meaning; [43, 44]). This atypical processing did not affect the accuracy of their responses, although the response times were longer for older adults than for young people. Older adults also showed a greater modulation of the amplitude in relation to the WM load than young people. Small amplitudes of the P600a and P600b components were observed in the high load condition of the MT, a fact that was not observed in the younger participants. These findings may be consistent with the idea of neural efficiency [9, 45] or the idea that older adults may show a less efficient neural response accompanied by compensation.

The objective of the present ERP study was to analyze the effect of CR on the performance of young adults. In the same sense as in the studies with older adults, we intend to see the CR effect by dividing the sample of participants into high and low CR groups. We hypothesize that young people with high CR will show better behavioral performance and that their brain electrical response pattern will reflect earlier processing and will be less vulnerable to the complexity of the task compared to the participants with low CR. We think that this beneficial effect of CR would be observed in young people facing two levels of WM load while they are performing tasks involving fluid intelligence (Sternberg WM task) and crystalized intelligence (Reading sentence task).

## Experiment 1

This experiment aimed to analyze specifically whether high levels of CR and two levels of memory load modulate the P300 amplitude and the behavioral performance of young adults during a Sternberg task. Given the evidence on greater P300 amplitude modulation and poor behavioral performance when participants have low CR [9], we expect to observe a lower percentage of correct answers and longer reaction times during a Sternberg task with two levels of WM load in a group of young people with low CR than in a second group with high CR. This pattern of differences would be more evident for the higher memory load condition. Likewise, the high CR group would show larger P300 amplitudes in both WM load levels than the low CR group. This latter group would also show a greater modulation of the P300 amplitude depending on the WM load; specifically, a smaller P300 amplitude is expected when the WM load increases. In contrast, participants with high CR would not show this pattern of amplitude modulation between WM loads because these conditions would not have a significant processing cost for this group.

### Materials and Methods

#### Participants

The sample consisted of 40 young Mexican adults, 18 women and 22 men. All participants were right-handed, and their ages ranged between 20 and 30 years (mean age = 24.4 years, SD = 3.04). All participants had normal or corrected vision and did not report any history of neurological or psychiatric problems, illegal drug use or frequent consumption of medical drugs and alcohol. In the case of women, the assessment was performed avoiding menstrual days [46].

Participants were grouped into two levels of CR (high and low) according to four proxy measures of cognitive reserve: a) **years of formal education** (high: ≥12 years, low: < 12), b) **occupational level** (high: levels 4 and 5, low: levels 1 and 2) from of the competencies of the National Classification System of Occupations [47] (level 1: jobs that require relatively easy tasks that could involve physical strength and endurance; level 2: occupations that require the use of specialized equipment or vehicles; level 3: jobs that demand more complex activities in which an intermediate level of knowledge in a more specialized field is needed; level 4: occupations involving decision making, problem solving, leadership skills, a specialized degree, an expert level of knowledge and wide experience in the field), c) **the verbal comprehension index** (high: VCI > 100, low: VCI ≤ 100,) from the WAIS-IV intelligence scale [48] and d) **the total score of a CR questionnaire** (high: > 90, low: < 70), which assess everyday activities.

Each participant received a score of 1 or 2 if his/her proxy scores fell into the low or high categories, respectively. The total score for each subject was obtained by summing these four proxy measures. Participants who obtained a total score of 7 or 8 were included in the high CR group, while those who obtained a total score of 5 or less were included in the low CR group. Subjects with a total score of 6 were excluded. Finally, 22 participants were placed in the high CR group and 18 in the low CR group. There were no significant differences regarding gender between the groups (high CR: 40% women, low CR: 50% women, X^2^ = 0.331, p = 0.56). Table 1 summarizes the sociodemographic features of the groups.

**Table 1.** Demographic data, WAIS scores and Cognitive Reserve questionnaire total score. Means and standard deviations.

|  | High CR | Low CR |  | 95%<br>Confidence<br>Interval |
| --- | --- | --- | --- | --- |
|  | N = 22<br>(female = 9) | N = 18<br>(female = 9) | t(38) |  |
|  | Mean (SD) | Mean (SD) |  |  |
| Age (years) | 23.18 (2.13) | 26.00 (3.33) | -3.11** a | -4.67 : -0.96 |
| Years of Schooling | 15.59 (1.71) | 10.33 (2.95) | 6.70*** | 3.64 : 6.87 |
| Occupation; mode (range) | 1 (3) | 1(1) | Z = -1.45 b | U = 148.5 |
| WAIS-Similarities | 11.59 (1.33) | 8.67 (2.20) | 5.20*** | 1.80 : 4.06 |
| WAIS-Vocabulary | 12.27 (2.10) | 8.94 (1.76) | 5.36*** | 2.07 : 4.59 |
| WAIS-Information | 11.77 (2.49) | 6.33 (2.20) | 7.25*** | 3.92 : 6.96 |
| WAIS-VCI | 110.68 (9.80) | 89.17 (8.47) | 7.33*** | 15.06 : 27.45 |
| Total CR questionnaire<br>score | 109.09 (37.97) | 123.39 (53.92) | -0.98 | -43.76 : 15.17 |

SD: Standard Deviation; CR: Cognitive Reserve; VCI: Verbal Comprehension
Index; a: $t(27.8)$ ; b: $p = 0.15$ , Mean Rank: High CR = 22.75, Low CR = 17.75. \*\* $p < 0.01$ ; \*\*\* $p < 0.001$

Informed consent was obtained from all participants. The project was approved by the ethics committee of the Institute of Neurobiology of the Universidad Nacional Autónoma de México (Ethical Application Ref: INEU / SA / CB / 109), according to the Declaration of Helsinki.

#### CR assessment

Similar to questions in the instruments proposed by others [49–51], we elaborated 26 questions with a Likert-type response assessing participant’s activities in four periods of their life (1 [18 – 21 years old], 2 [22 – 25 years old], 3 [26 – 29 years old] and 4 [30 – 33 years old]). These questions were intended to assess everyday activities (i.e., social, cognitive and physical activities). The response options were framed in terms of the frequencies of these activities: *0) Never, 1) A few times a year, 2) A few times a month, 3) A few days a week and 4) Almost every day or every day*. Each participant give one or more answers to each item depending on their age (number of periods);the older the participant was, the more periods of life they had completed. The total score for each person was obtained by directly adding the scores given from the items across the periods. A higher scores indicates a higher frequency of different activities and a higher cognitive reserve. The possible maximum score per item was 16, and the maximum total score was 416.

The reliability and validity of our CR questionnaire was assessed by applying it to 193 Mexican young people (73 women and 120 men) between 20 and 30 years old. The internal consistency of the 26 items was acceptable (Cronbach’s α = 0.950). Pearson correlation analysis showed significant positive correlations between the total score of the questionnaire and the educational level (r = 0.449, p < 0.001), occupational level (r = 0.264, p < 0.01), and the VCI (r = 0.273, p < 0.01) of these participants.

#### Stimuli

One hundred eighty digit-sets divided into two levels of memory load were built. The memory sets were formed by four digits (1 to 9). The low memory load sets consisted of only one digit repeated four times; in contrast, the high memory load sets consisted of four different digits randomly sorted, avoiding strings of consecutive numbers, either in ascending or descending order. The digits were presented in white (Arial, size 12) on a black background. The probe stimulus was a single digit presented at the center of the monitor. Trials were presented randomly during the trial sequence and divided into two blocks with 45 trials each (90 for each level of difficulty).

#### The Sternberg task

The WM task trials were a modified version of the Sternberg verbal work memory paradigm and began with the presentation of an asterisk for 1000 ms at the center of the screen (fixation point); this was followed by an interstimulus interval of 500 ms, as shown in Fig 1. Subsequently, a memory set of four digits was presented for 1000 ms. A retention time of 1000 ms was then initiated where the screen turned black at the end of the presentation of the set. Next, a probe digit was presented for 1000 ms, and the participants were instructed to respond by pressing one button on a response box if the probe digit was included in the memory set previously presented (positive probe) or the other button if the probe digit had not appeared (negative probe). The use of the response buttons was counterbalanced across participants. Memory set trials (high and low WM load) and probe stimuli (positive or negative) were randomly presented in the trial sequence. The participants were instructed to answer as quickly as they could while avoiding making mistakes. Participants were not provided feedback on their performance. The task lasted approximately 20 min.

**Fig 1.**
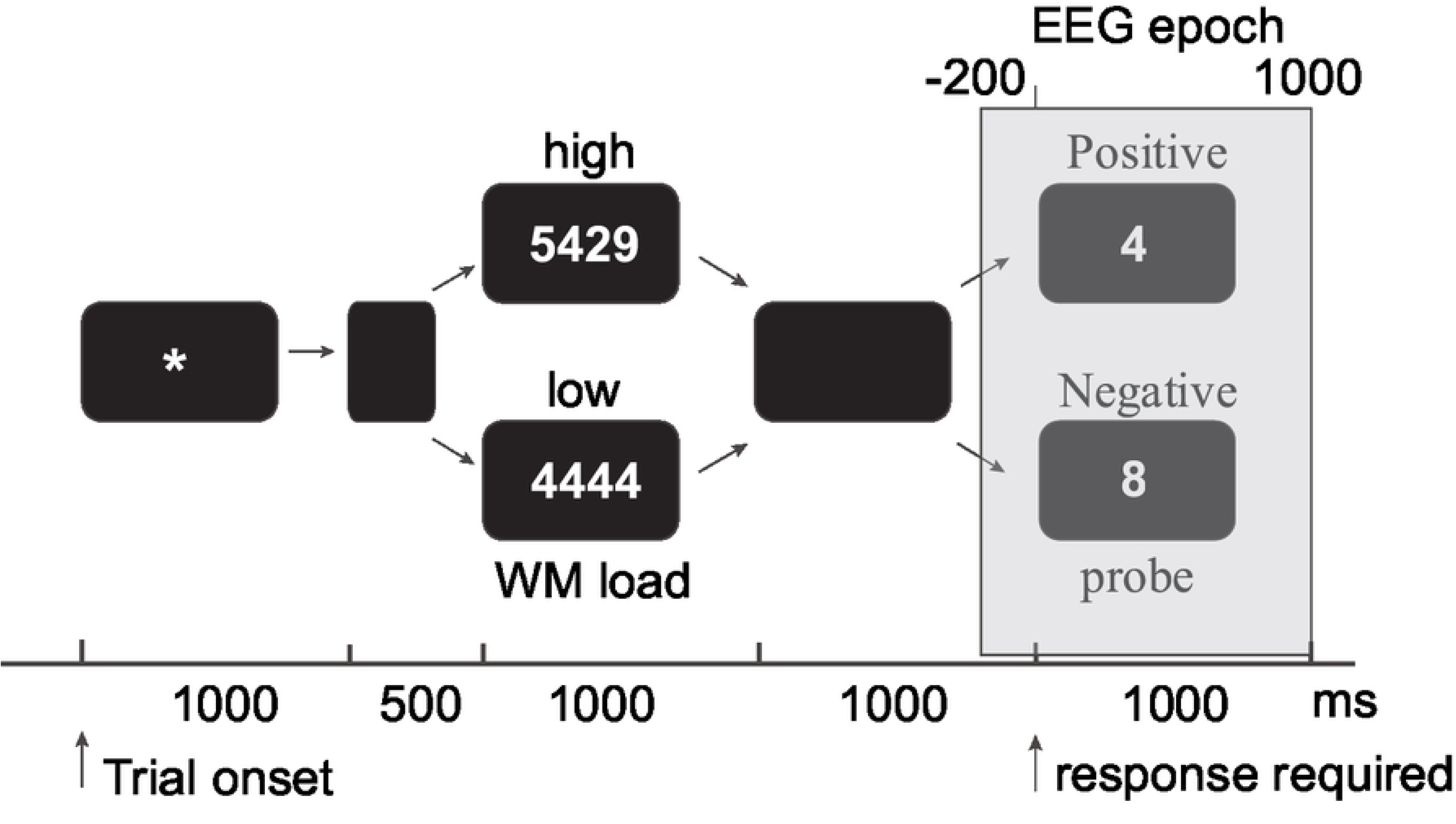
Time flowchart of Sternberg’s working memory task. Examples of high and low WM load condition memory sets and probes that do or do not belong to the set (positive and negative probes). The gray box represents a window time-locked to the continuous EEG.

The stimuli were delivered by the STIM2 software (NeuroScan, CompuMedics, Charlotte, NC, USA) through a PC using a 17” monitor. Participants sat 70 cm from the screen in a room with the light off.

#### ERP acquisition and analysis

The electroencephalogram (EEG) was recorded using a SynAmps system (NeuroScan, CompuMedics) and collected through 32 silver electrodes embedded in an elastic cap (ElectroCap International, Inc., Eaton, OH, United States) referenced to the right earlobe (A2). The left earlobe was recorded independently. The bandwidth of the amplifiers was set to 0.1-100 Hz, and the signal was digitized at a 500 Hz sampling rate. Impedances were kept below 5 kΩ. Two electrodes placed on the external canthus and superciliary arch of the left eye were used to record the electrooculogram (EOG).

The EEG recordings were rereferenced offline using the average of the earlobe signals (A1-A2). The continuous EEG recording was epoched from 200 ms prestimulus to 1000 ms poststimulus. The ERP waveform was baseline corrected, and drift was removed using the linear detrend tool from the NeuroScan 4.5 software (CompuMedics). All EEG epochs were visually inspected, and manual rejection of segments was performed. Segments corresponding to incorrect responses were rejected. The artifact-free segments from each participant were averaged for each experimental condition. Approximately 23 segments were used for the average of the ERPs for both groups in all task conditions. No significant differences were observed regarding the number of artifact-free EEG segments across experimental conditions. From the total EEG segments of the high CR group, 58.5% of the high WM load condition segments and 61.5% of the low WM load condition segments were retained. For the low CR group, 56% of the high WM load condition segments and 57% of the low WM load condition segments were retained.

#### Statistical analysis of the behavioral data

Analysis of the behavioral data was carried out using the commercial software SPSS 20. Two three-way ANOVAs were performed on the percentage of correct answers and the response times. WM load (high and low) and Probe type (Positive and Negative) were used as within-subjects factors and CR level (high and low) was used a between-subjects factor. Honest significant difference (HSD) post hoc pairwise tests were performed for multiple comparisons. The percentage of correct answers for both tasks was transformed using the function arcsine[square root(percentage / 100)] to ensure the normal distribution of the data.

#### Statistical analysis of the ERP data

For the ERP latency data, two three-way ANOVAs were separately performed on the P300 latencies by Probe type (Positive and Negative) with WM load (high and low) and Electrode site (Fz, Cz, Pz) as within-subjects factors, and CR level (high and low) as a between-subjects factor. Honest significant difference (HSD) post hoc pairwise tests were performed for multiple comparisons.

For the analysis of ERP amplitude, the nonparametric permutation test was performed. Since there is a multiplicity of comparisons in the analysis of the ERP data, there is a high probability of increasing the type I error, so the use of multivariate nonparametric permutation tests has been recommended [52]. This method is a distribution-free test that builds an empirical probability distribution computing multiple statistical tests. This statistical tool is included in the eLORETA software (exact low-resolution brain electromagnetic tomography; [53]). The statistical analysis was performed using 10,000 permutations. t _max_ (0.05 p-alpha) and its global p-value were reported. This statistic represents the significance when all the electrodes sites and/or all the points in times are considered. Significant t-marginal values (all p < 0.05) corresponding to specific differences at each electrode site were represented in color maps. The false discovery rate (FDR) method was used to correct the global p-alpha for multiple comparisons between experimental conditions using nonparametric analysis with permutations. We used the Benjamini-Hochberg method [54] to identify those p-values associated with t-max global values that remain significant.

### Results

#### Behavioral

The percentage of correct answers and the average response time are shown in Fig 2. A WM load by CR group interaction was found (F(1, 38) = 7.66, p = .009, η_p_^2^ = 1.7) for the percentage of correct answers. The post hoc tests showed that both groups (high CR: MD _HSD_ = −0.11, p < 0.001; low CR: MD _HSD_= −0.20, p < 0.001) had a larger percentage of correct answers for the low WM load than for the high WM load, but this difference was greater for participants with a low CR.

**Fig 2.**
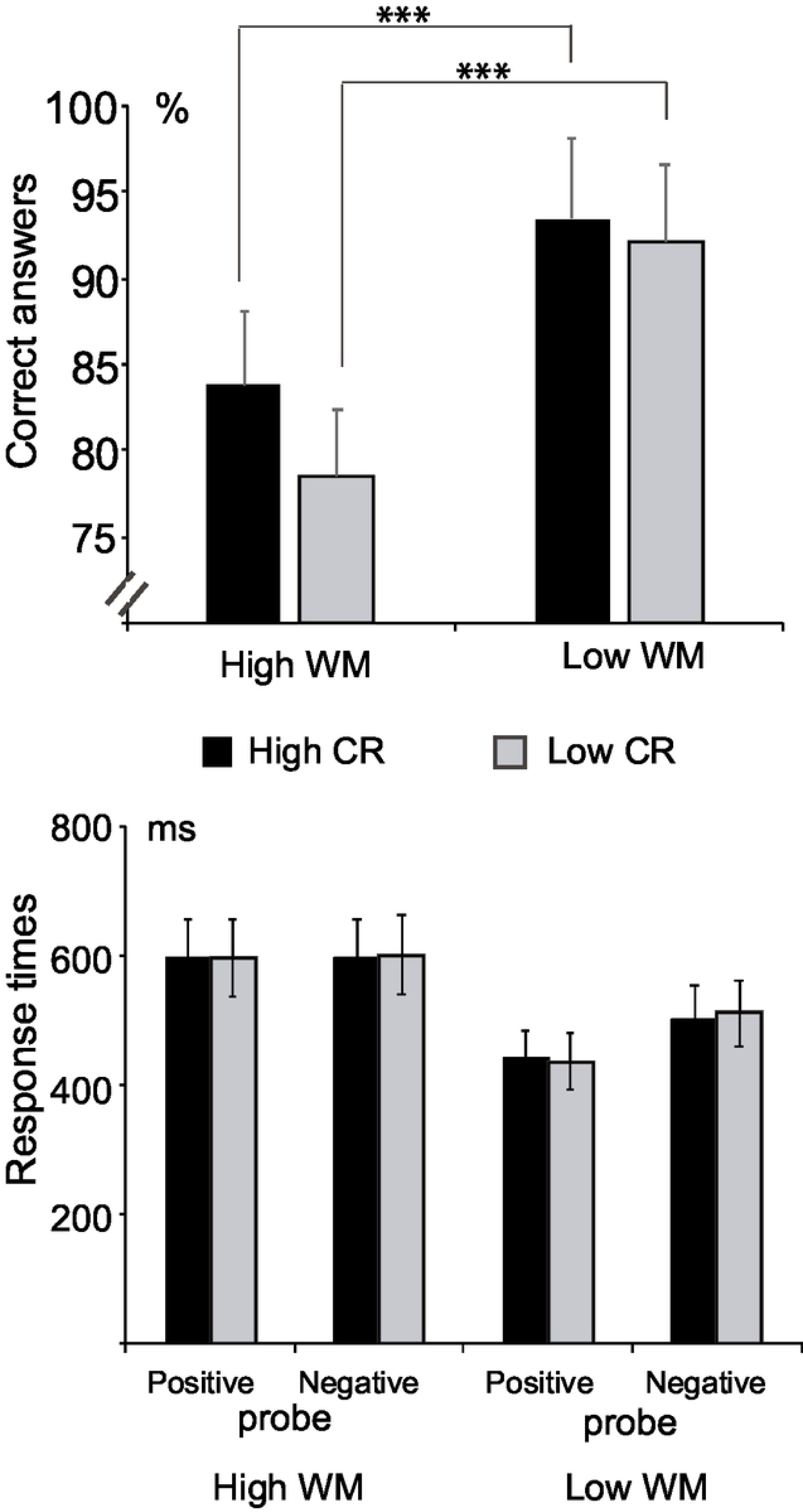
Means and standard deviations of the percent of correct answers (top) and response times (bottom) for high and low working memory (WM) sets. Note that both groups displayed significantly lower percentages of correct answers for the high WM load than for the low WM load condition. This effect was greater for the group with low CR (represented in gray bars). *** p < 0.001

There were no significant differences between the groups with respect to the response time (main effect of Group: F < 1), the WM load by CR group interaction (F < 1) or the Probe type by CR group interaction (F < 1).

#### ERP time windows

To delimit the time windows where the ERP amplitude differed between probe types (positive and negative) throughout the whole epoch (−200 to 1000 ms) and across all electrodes over the scalp, a nonparametric permutation test was performed. Two independent analyses were separately performed for each CR group, one for each probe type.

Fig 3a shows the grand averages of each group for each memory set condition. In both groups and both probe types, significant differences between WM loads were found in the interval from 270 ms to 450 ms (high CR group: positive probes [t _max_ (.05) = −5.359, global p < .0001]; negative probes [t _max_ (.05) = −5.464, global p < .0001]; low CR group: positive probes [t _max_ (.05) = −5.355, global p < .001]; negative probes [t _max_ (.05) = −5.580, global p < .0001]). Specifically, within this time window, the positive wave had a significantly larger amplitude for the low WM load than for the high WM load condition. Taking into account the latency of occurrence, the polarity, and the correspondence of this component with this cognitive task, this significant effect can be considered as the P300 component.

#### ERP latency

There were greater P300 latencies for the high WM load than the low WM load condition for the positive probes (F(1,38) = 26.9, p < 0.0001, η_p_^2^ = 0.41) and for the negative probes (F(1, 38) = 5.75, p = 0.021, η_p_^2^ = 0.13). The WM load by Electrode site by Group interaction was not significant for the positive probes (F(2, 76) = 2.65, p = 0.083, η_p_^2^ = 0.07, ε _G-G_ = 0.91), but the interaction was significant for the negative probes (F(2, 76) = 3.21, p = 0.049, η_p_^2^ = 0.08, ε _G-G_ = 0.95). Pairwise multiple comparisons indicated that participants with high CR showed slightly greater P300 latency to high WM than low WM loads at Cz (MD _HSD_ = 31.05, p = 0.048), while participants with low CR displayed a more-significantly greater P300 latency to high WM than low WMs load at Pz (MD _HSD_ = 49.56 p = 0.001)

#### ERP amplitude

Fig 3b shows statistical maps that represent the comparisons between WM loads using the mean amplitudes at the P300 window for each Group and Probe type. In both groups, there was a larger P300 amplitude for low WM loads than for high WM loads, both for positive and negative probes, over all electrode sites. In the high CR group, a larger P300 amplitude for the low WM load than the high WM load condition was observed for positive probes (t _max_ (.01) = 3.57, global p = 0.0002) over all electrode sites (all p < 0.001), but mainly over C4, P3, T3 and T5 (p < 0.0001). In this group, there was also a larger P300 amplitude for low WM loads than for high WM loads for negative probes (t _max_ (.01) = 3.39, global p = 0.0002) over all electrode sites (all p < 0.001), but mainly over Ft8, T3, T4, Tp7 and Tp8 (p < 0.0001). Similarly, these differences were observed in the low CR group for positive probes (t _max_ (.01) = 3.64, global p = 0.0002) over all electrode sites (all p < 0.001), but mainly over P3 and P4 (p < 0.0001), and for negative probes (t _max_ (.01) = 3.26, global p = 0.0002) over all electrode sites (all p < 0.001), but mainly over C4, P4, Cpz, Cz, and Cp4 (p < 0.0001).

A comparison between WM loads by Groups was performed for each type of probe (H0: A1-A2 = B1-B2, where A and B are the high and low CR groups, respectively, and 1 and 2 are high and low WM loads, respectively). There were no significant differences for positive probes (t _max_ (0.05) = 2.636, global p= 0.382) or for negative probes (t _max_ (0.05) = 2.629, global p = 0.520). However, as can be noted in Fig 3b, the distribution of the differences between WM loads seemed to be wider for the low CR group than for the high CR group when negative probes were used. Furthermore, these topographic differences in the P300 amplitude seemed to have a wider distribution in the negative than in the positive probe conditions for the low CR group than for the high CR group.

**Fig 3.**
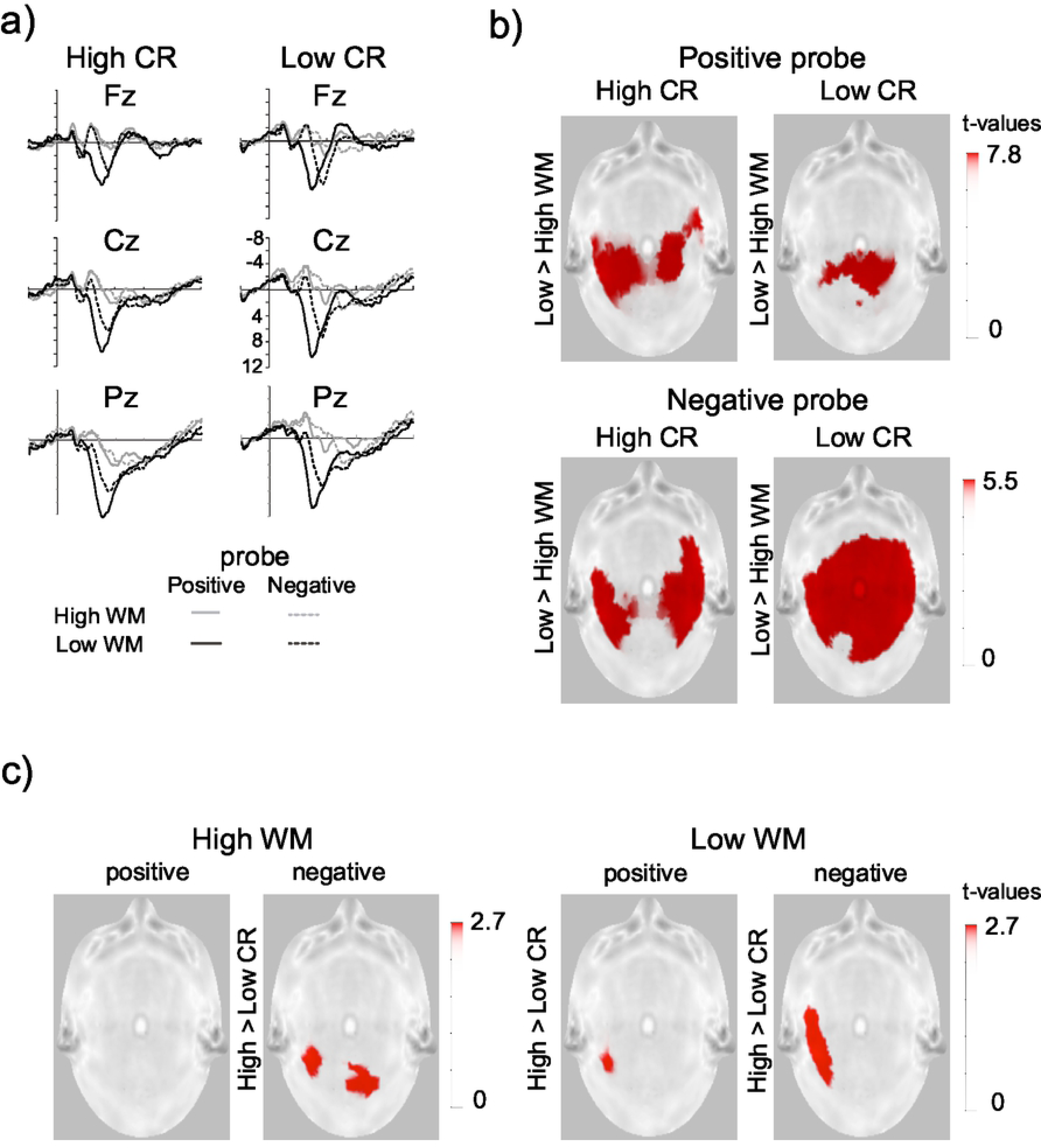
Event-related potentials to the Sternberg task. a) Grand averages of the high cognitive reserve (CR) and the low CR groups at the midline electrodes. Negative is plotted up. Gray lines represent the ERPs to high WM loads and black ones to low WM loads. Solid lines represent positive probes, and dotted lines represent negative probes; b) statistical maps of the amplitude change of the P300 component by WM load for each group and type of probe. Red represents t-values where the P300 amplitude change was significant (low WM load > high WM load); c) statistical maps of the P300 effect for positive and negative probes for the two WM load conditions (high and low). A significantly greater P300 effect is represented with red dots.

In another series of analyses, four independent samples t-tests were performed. Fig 3c shows the comparison between the two groups (high and low CR) for each of the four experimental conditions. For high WM load and positive probes, the high CR group showed a greater amplitude of the P300 component than the low CR group (t _max_ (.05) = 2.679, global p = .0064, FDR _H-B_ p = .0128) over the posterior recording sites (all p < 0.05). In this same WM load condition, but for negative probes, there were no differences between groups (t _max_ (.05) = 2.731, global p = .154, FDR _H-B_ p = .1540). In the low WM load condition, there were P300 amplitude differences between the groups for both types of probes. For the positive probe, the amplitude of the P300 component was larger in the high CR group than in the low CR group (t _max_ (.05) = 2.750, global p = .0200, FDR _H-B_ p = .0266) over T5, and for the negative probe, the amplitude of the P300 component was also larger in the high CR group than in the low CR group (t _max_ (.05) = 2.732, global p = .0054, FDR _H-B_ p = .0216), here over T3 and Tp7.

### Discussion

It was expected that the group of young people with high CR would show greater efficient and faster responses than the group with low CR in a Sternberg-type task. Additionally, it was expected that those with high CR would show smaller changes in their behavioral performance between different levels of WM load than those with low CR. In accordance with our hypotheses, the participants with high CR showed smaller differences as the WM load increased in the percentage of correct answers than those with low CR. This finding supports the idea that a higher CR is associated with efficient neural networks operating to perform a cognitive task and is consistent with previous studies in older [6,9,55–57] and young adults [9].

In contrast to previous results in young adults using a Sternberg-like task [6], where significant correlations between CR composite scores and response times were observed, but consistent with Speer & Soldan [9], our groups of young participants did not differ in response times. Our findings can be interpreted as the use of different strategies to solve the task. Persons with high CR could be more precise while sacrificing response time, while those with low CR responded as slowly as the others but more erratically.

Considering the idea of neural efficiency [45], we also hypothesized that the effect of CR could be observed in changes in P300 amplitude with respect to WM load. According to Speer & Soldan [9], this idea could be considered a measurement of neural inefficiency. In their study, they correlated the mean change in the amplitude of the P300 component as a function of WM load with the CR composite and with behavioral performance. To calculate the mean amplitude change, they averaged the amplitudes across the P300 time window and across electrodes for each set size. Afterwards, they subtracted amplitude at set size 1 from the amplitude at set size 7 Then, similar to Gu et al.’s results [6], they found that individuals with higher CR composite scores showed less change in P300 amplitude with increasing task demands. Based on these findings, we expected that the participants with low CR should show greater modulations or changes in the P300 amplitude as the WM load increased (i.e., greater P300 amplitude changes in the low than in the high WM load condition).

Both groups of young adults displayed significantly larger amplitudes of P300 for low WM than for high WM loads (WM load-related changes). Although there was no evidence of a significant difference between groups with increasing WM load, the low CR group seemed to display a wider distribution of WM load-related changes than the high CR group to negative probes. Why was the previously reported pattern not observed here? A possible explanation is that the previous studies computed a composite of P300 amplitudes from all electrode sites, from all time points within the P300 time window and for both the positive and negative probe conditions; then, this composite was correlated to the CR composite score. This procedure could hide some topographical differences between the groups by the type of probe. Our results could indicate that participants with low CR, on the one hand, have an inefficient neural network, which was shown by the P300 amplitude changes as the WM load increased for the negative probes, and on the other hand, were as efficient as participants with high CR when positive probes were used, perhaps because the processing of negative probes requires a complete search of the set, implying more cognitive demand. This could be supported by our latency findings and by the results of previous studies [6, 9]. The low CR group displayed significantly longer P300 peak latencies to high WM than to low WM loads, and this difference was more marked in this group than in the high CR group for the negative probes.

A higher cognitive demand was also observed globally across experimental conditions, even though there were no robust behavioral differences between the groups. The participants with low CR showed a significantly smaller P300 amplitude than those participants with high CR for negative probes at high and low WM loads and for positive probes at low WM loads. These findings are supported by previous P300 ERP studies [33, 34] that showed that smaller amplitudes of the P300 component are associated with greater difficulty of processing, while larger amplitudes are related to less difficulty in performing the task while maintaining a constant level of performance.

## Experiment 2

In this experiment, we analyzed the effect of CR on the performance of young people, wherein grammatical gender agreement is processed during a grammatical judgment task of sentences. There is scarce evidence to indicate different cognitive performance between subjects with high and low CR in sentence processing during reading. In healthy older adults, in whom cognitive decline is more likely to manifest, there is a more efficient response when CR is greater (habitual reading activities) [40]. It is feasible to think that CR, considering mainly the educational level as a proxy measurement, has a positive effect on the behavioral strategies and on the variety of brain mechanisms that make solving problems effective [58]. Thus, we expected that during the reading of sentences, the processing of gender agreement would lead to a higher processing cost when there is a higher WM load than when there is not. The high CR group would show better behavioral results than the low CR group when processing sentences with a low WM load. We also think that young people with low CR will base their response on a strategy where the analysis of semantic information is preferred to resolve the gender agreement, because this would result in a smaller amplitude of the LAN effect than that of subjects with high CR. Participants with low CR will also show a larger amplitude of the P600a effect because they will be forced to compensate for the poor processing of the previous stage. In the final stage of the agreement processing, during the P600b window, where the generalized mapping of the sentence is made and the semantic and syntactic information converges, there will probably be no differences between the two groups.

### Method

#### Participants

Participants in Experiment 2 and their classification into the two levels of CR groups were the same as in Experiment 1.

#### Stimuli

The task consisted of 160 experimental sentences with seven words each. Adjective-noun grammatical gender agreement (agree and disagree) and WM load (high and low) were manipulated. Table 2 shows examples of the sentences presented to the participants. Forty sentences correspond to the agree / low WM load condition, where the grammatical gender of the main noun in the sentences agrees with the gender of the adjective. To build 40 disagree sentences (disagree / low WM load condition), the grammatical gender of the qualifying adjective that modifies the main noun of the sentence was manipulated. To build 40 agree sentences with a high WM load condition (agree / high WM load condition), four words were embedded between the noun and the adjective. Forty disagree sentences with a high WM load were also included. Additionally, sixty filler sentences were added. In these sentences, the number agreement between the noun and adjective was manipulated. Fifty percent of the filler sentences were agree sentences, and the remaining were disagree sentences.

**Table 2.** Examples of sentences for each experimental condition.

| WM Load | Agreement | Example |
| --- | --- | --- |
| Low | Agree | La **casa**<sub>female</sub> **roja**<sub>female</sub> está en la colina |
|  |  | The **red**<sub>female</sub> **house**<sub>female</sub> is on the hill |
|  | Disagree | El **edificio**<sub>male</sub> **roja**<sub>female</sub> está en la colina |
|  |  | The **red**<sub>female</sub> **building** is on the hill |
| High | Agree | La **casa**<sub>female</sub> que está allá es **roja**<sub>female</sub> |
|  |  | The **house**<sub>female</sub> over there is **red**<sub>female</sub> |
|  | Disagree | El **edificio**<sub>male</sub> que está allá es **roja**<sub>female</sub> |
|  |  | The **building**<sub>male</sub> over there is **red**<sub>female</sub> |
| Filler | sentences | Las casas<sub>plural</sub> **amarilla**<sub>singular</sub> están en la colina |
|  |  | The **yellow**<sub>singular</sub> houses<sub>plural</sub> are on the hill |
|  |  | Las casas<sub>plural</sub> que están allá son **amarilla**<sub>singular</sub> |
|  |  | The houses<sub>plural</sub> over there are **yellow**<sub>singular</sub> |

Note: Nouns and adjectives are typed in bold fonts.

#### Grammatical judgment task (reading sentences)

Each sentence began with the presentation of an asterisk for a duration of 300 ms at the center of the monitor, and then the sentence appeared word by word as shown in Fig 4. Each word was presented for 300 ms with an interstimulus interval of 300 ms. At the end of the sentence, two question marks were presented for 500 ms, and participants had 1500 ms to indicate whether the sentence did or did not have correct Spanish grammar using a response box. Pressing one button indicated that the sentence was correct, and the other indicated that it was wrong. The use of buttons was counterbalanced among the participants. The task lasted approximately 40 min, and as in Experiment 1, the stimuli were delivered by STIM2 software (Compumedic, NeuroScan).

**Fig 4.**
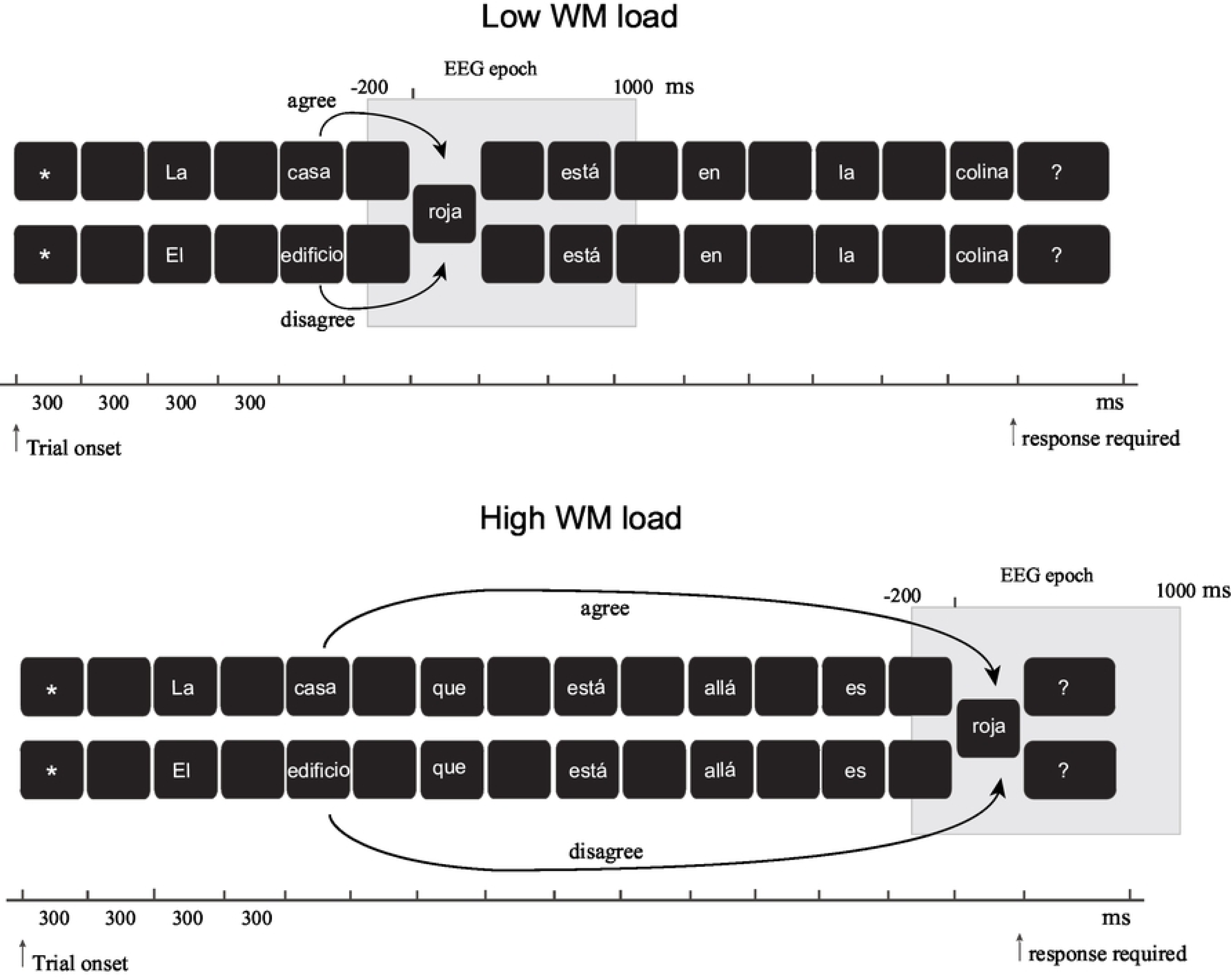
Time flowchart of the sentence presentation (word by word) for the grammatical judgment task (reading sentences). Two example sentences (agree and disagree) in the low WM load condition, where the arrows display the proximity between noun and adjective. Two example sentences in the high WM load condition, where the noun is separated from the adjective by four words. The gray boxes represent the moment where the EEG epoch is taken for the analysis.

#### ERP acquisition and analysis

This experiment followed the same procedure as Experiment 1 regarding EEG acquisition and preprocessing of the EEG signal. For this grammatical judgment task, the continuous EEG signals were synchronized to adjective onset (−200 to 1000 ms), and the EEGs were segmented according to the different experimental conditions (agree / low WM load, disagree / low WM load, agree / high WM load, disagree / high WM load). The number of segments used to obtain the ERPs in each experimental condition for both groups was approximately 23. The percentage of artifact-free segments was 59.8% for low WM loads and 59.1% for high WM loads in the high CR group and 53.4% and 51.3%, respectively, in the low CR group.

#### Statistical analysis of the behavioral data

The analysis of the behavioral data was performed similarly to that in Experiment 1. Two three-way ANOVAs were performed on the transformed percentage of correct answers and the response times. WM load (high and low) and Gender agreement (agree and disagree) were used as within-subjects factors, and CR level (high and low) was used as a between-subjects factor. HSD post hoc tests were performed for multiple comparisons.

A multivariate nonparametric permutation test with 10,000 permutations was used to analyze the ERP amplitude data. The FDR method and Benjamini-Hochberg correction were used to correct the global p-alpha for multiple comparisons between experimental conditions.

We proceeded first with comparing the agree versus disagree conditions for each level of WM load and the two groups of participants across all electrode sites and points of time (whole time epoch). The significant differences between conditions allowed us to define the time windows for posterior analyses, which would correspond with the ERP components associated with gender agreement processing. Second, ERP difference waves were obtained by subtracting the amplitude values of the whole-epoch ERP to the agree from those to the disagree condition for each participant. Using independent samples t-tests based on the permutation technique, a series of analyses was carried out using the mean amplitude value of the difference wave (i.e., disagree minus agree) for comparing between CR groups for each level of WM load at each time window determined by multivariate permutation analysis. Third, we compared the mean amplitude values at each time window between WM loads for each CR group (high and low).

### Results

#### Behavioral

Fig 5 shows the percentages of correct answers and the average response times for gender agreement processing in the grammatical judgment task. The high CR group had a higher percentage of correct answers in all the conditions of the grammatical judgment task than the low CR group (main effect of Group: F(1,38) = 13.18, p = 0.001, η_p_^2^ = 0.258).

Regarding response time, there was a significant main effect of CR group (F(1, 38) = 14.85, p < 0.001, η_p_^2^ = 0.281), which means that young participants with high CR responded faster than those with low CR independent of the experimental condition. There was also a significant Gender agreement by CR Group interaction (F(1, 38) = 7.88, p = 0.008, η_p_^2^ = 0.172) and Gender agreement by WM load by CR Group interaction (F(1,38) = 7.008, p = 0.012, η_p_^2^ = 0.156). The post hoc tests of the last interaction indicated that the participants with high CR responded faster than those with low CR in the low WM load condition for the agree (MD _HSD_ = −99.72, p = 0.002) and disagree conditions (MD _HSD_ = −101.24, p < 0.001); they also responded faster in the high WM load condition for the agree (MD _HSD_ = −80.05, p = 0.01) and disagree conditions (MD _HSD_ = −145.13, p < 0.001), as shown in Fig 5.

**Fig 5.**
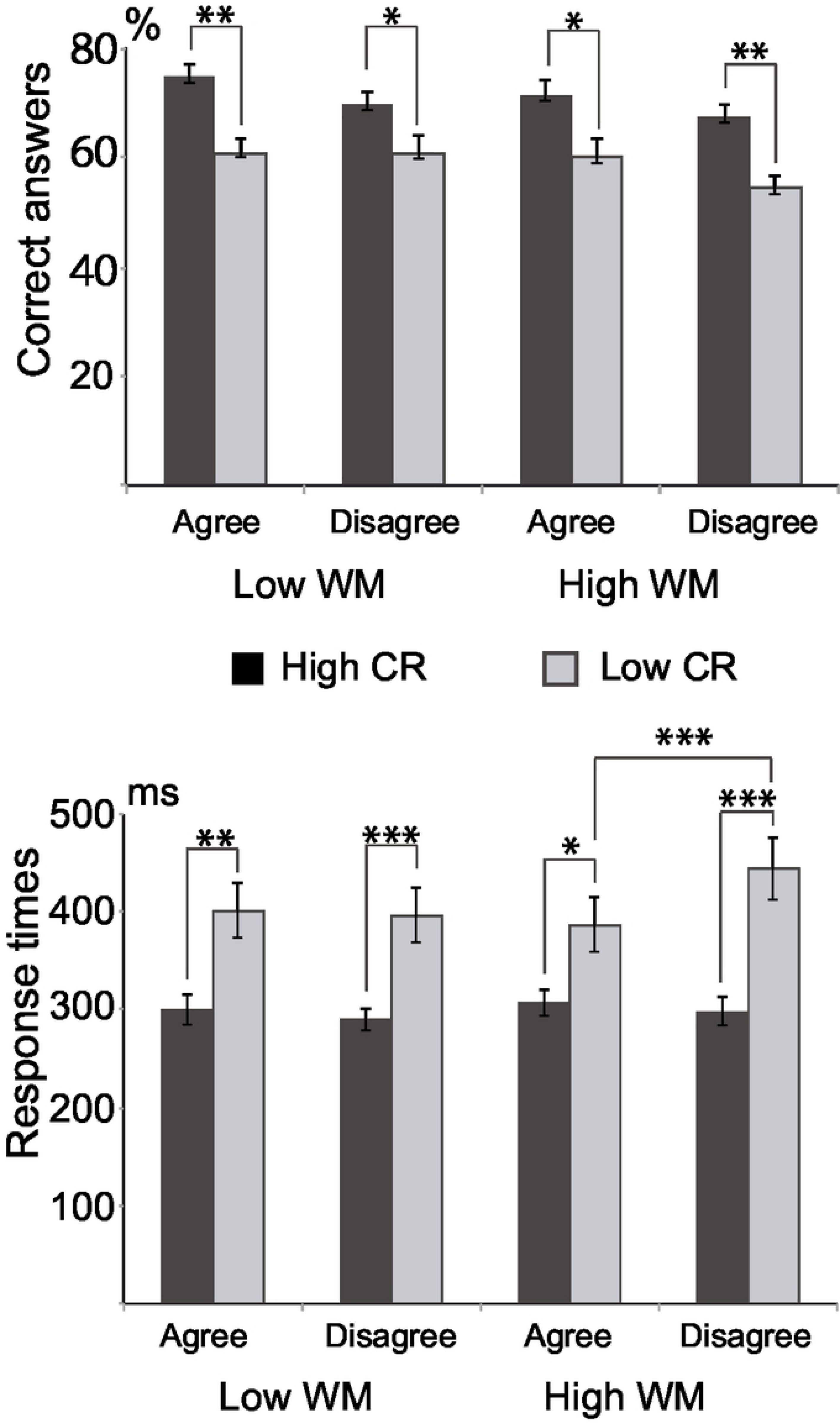
Behavioral results in the grammatical judgment task. Percentages of correct answers (top) and average response times (bottom). The group of participants with high CR showed a higher percentage of correct answers and faster response times than those with low CR. Note that young adults with low CR show longer response times to disagree than agree sentences at the high WM load condition. * p < 0.05, ** p < 0.01, *** p < 0.001

#### ERP time windows

To determine the time widows where significant differences between agree and disagree conditions were found, a series of nonparametric permutation statistical tests was performed for each CR group and WM load.

Fig 6a shows the grand averages of the CR groups for each experimental condition. For each level of WM load, both groups showed significant differences between agreement conditions at three consecutive time windows, which correspond to the occurrence of LAN (330 – 380 ms), P600a (520 – 660 ms) and P600b (700 – 800 ms) effects (high CR low WM load: t _max_ (.05) = −5.464, global p = 0.003; high CR WM load: t _max_ (.05) = −5.538, global p = 0.0004; low CR low WM load: t _max_ (.05) = −4.919, global p = 0.028; low CR high WM load: t _max_ (.05) = −5.47, global p = 0.0080).

#### ERP amplitude

Difference waves were obtained by subtracting the amplitude values of the whole-epoch ERP for the disagree condition from those for the agree condition for each participant. Using independent samples t-tests based on the permutation technique, a series of analyses was carried out comparing the CR groups using the mean amplitude value of the difference wave (i.e., disagree minus agree) at each time window corresponding to LAN, P600a and P600b.

First, for each WM load condition, the difference wave of the high CR group was compared to the difference wave of the low CR group. As shown in Fig 6b, f or high WM loads, the participants with high CR had a greater LAN effect (t _max_ (.05) = −2.844, global p = 0.0132, FDR H-B p = 0.0395) over F7, Fc4, C3, C4, T3, T5, Cp3, Cp4, Cz, Cpz and Tp7. They also showed a greater amplitude of the P600a effect than those in the low CR group (t _max_ (.05) = 2.594, global p = .0498, FDR H-B p = 0.074) over Cp4 and Cpz. The P600b effect was not significantly different between the groups (t _max_ (.05) = 2.566, global p = 0.2193, FDR H-B p = 0.2193). At low WM loads, there were no significant differences between groups in terms of the LAN (t _max_ (.05) = 2.673, global p = 0.689), P600a (t _max_ (.05) = 2.710, global p = 0.2968) or P600b effect (t _max_ (.05) = 2.783, global p = 0.1476).

**Fig 6.**
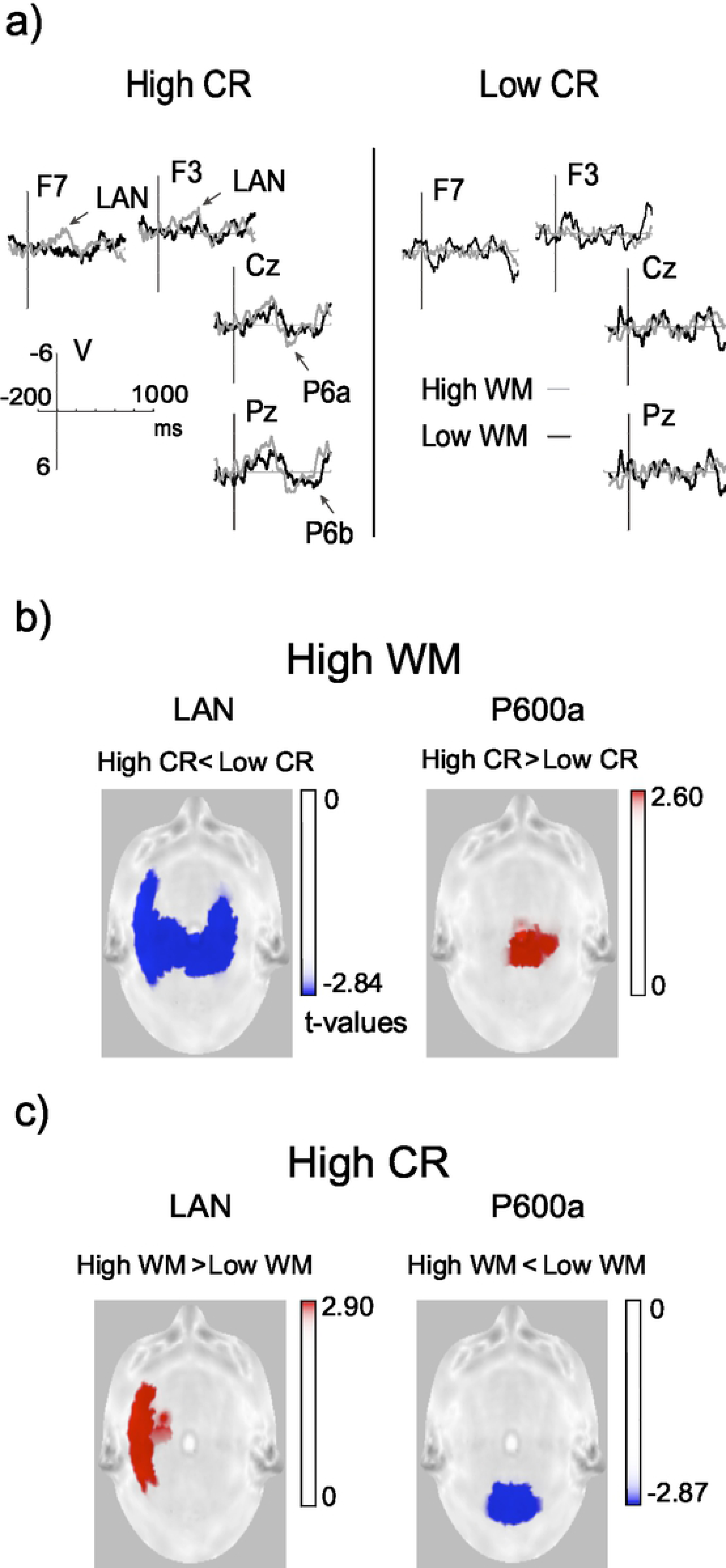
Statistical results of ERP amplitude analyses for the grammatical judgment task. a) Difference waveforms for the agreement effect (i.e., disagree minus agree condition) in the high and low CR groups. Negative is plotted up. The solid gray line represents the high WM load condition and black lines represent the low WM load condition. Note the left anterior negativity (LAN) effect at the left frontal electrodes (F3, F7), the P600a effect at the vertex electrode (Cz), and the P600b effect at the parietal site (Pz) in the high CR group; b) statistical maps of the agreement effect in the high WM load condition for the LAN and P600a ERP components; colored dots represent significant (p < 0.05) t-values in the comparison between groups with different levels of CR; c) statistical maps of significant t-values for the high CR group for LAN and P600a; comparisons between WM loads.

Second, the difference wave corresponding to the high WM load condition was compared to the difference wave to the low WM load condition for each group of participants separately. Fig 6c shows statistical maps representing such comparisons. Maps show that the high CR group showed a greater LAN effect for high WM than for low WM load; t _max_ (.05) = 2.859, global p = 0.0012, FDR _H-B_ p = 0.0036) over C3, F7, T3, Fc3, Tp7 and Ft7. This group also showed a smaller amplitude of the P600a effect for high WM than for low WM loads (t _max_ (.05) = −2.869, global p = 0.030, FDR _H-B_ p = 0.0450) over posterior sites. There were no differences between high and low levels of WM load at the P600b time window (t _max_ (.05) = −2.791, global p = 0.064, FDR _H-B_ p = 0.064). In contrast, the low CR group displayed no differences between the levels of WM load for the LAN effect (t _max_ (.05) = 2.738, global p = 0.2278), the P600a effect (t _max_ (.05) = −2.700, global p = 0.2189) or the P600b effect (t _max_ (.05) = −2.622, global p = 0.4349).

### Discussion

Young participants with high CR showed better performance than the group with low CR in terms of the percentage of correct answers and response times across experimental conditions in the grammatical judgment task. Regardless of WM load, but in accordance with our expectations, a higher cognitive reserve allows individuals to have more efficiently process grammatical gender agreement during sentence reading. This finding can be supported by previous studies showing a beneficial effect of CR on the reading abilities of healthy older adult performance [40, 57].

Regarding the brain response pattern, young people with high CR showed a greater amplitude of the LAN effect than the participants with low CR in the high WM load condition. This finding supports the idea that CR has a positive effect from the first stage of processing of gender agreement, as was suggested in a previous study, where ERP amplitudes of healthy older adults were compared with those of young adults processing gender agreement [41]. Older adults who showed age-related subtle signs of cognitive decline displayed unsuccessful morphosyntactic processing (reflected by the LAN effect). Thus, if a morphosyntactic analysis [43,44,59] that interacts with WM [60] is given at the first stage of the gender agreement process, then increasing the cognitive demand by increasing the WM load [61] results in a higher cost of processing and a decrease in the amplitude of the LAN effect in young people with low CR, which influences the later consecutive stages of processing. In this sense, at the early phase of P600, probably related to a processing stage in which the adjective is integrated into the sentence representation [43, 44], young people with low CR must try to compensate and complete the unsuccessful previous stage of processing. The low CR participants in this group probably showed this pattern mainly at high WM loads because they displayed a smaller amplitude of the P600a effect than those with high CR. Although this result does not fully support our hypotheses, it does suggest that during this stage, the cost of processing is much greater and the integration of the arguments is likely inadequate. This pattern of brain response agrees with the results observed when comparing older versus younger adults [41].

With regard to neural efficiency, the effect of low CR on gender agreement during sentence processing would be observed as a modulation of the amplitude that depends on WM load [9]. In contrast to this hypothesis, our results showed that the participants with high CR had greater amplitude changes as the WM load increased. This modulation was observed from the first steps of grammatical gender processing. However, even this finding could fit with the idea of neural inefficiency in young people with low CR, especially since such a pattern of brain responses, i.e., no amplitude changes with an increase in WM load, is associated with poor behavioral performance and evidence of much greater processing costs during the reading of sentences. In this sense, given that there were no amplitude differences in the P600b effect in any of the two groups when the two levels of WM load were compared, this finding could probably show that young people with low CR have to reach the last step of gender processing (using syntactic and semantic information) to carry out sentence reanalysis and thus employ more resources, similar to previous results where young and older adults were compared [41].

In summary, a low CR, associated with worse behavioral performance and a higher cost of processing in the grammatical gender agreement, can be related to a lower amplitude modulation of the brain response as the WM load increases. Thus, a higher CR can be associated with a more efficient brain response for sentence processing. An important limitation of this study was not having measured the reading ability of the participants. This variable could bias the results.

## General discussion

In the present study, behavioral performance and brain response were analyzed in two cognitive tasks involving fluid intelligence (working memory) and crystallized processes (grammatical gender agreement) between two groups of young people with different levels of CR. In agreement with our hypothesis, young people with high CR showed better behavioral performance in both tasks than young people with low CR. This improvement effect, probably induced by CR, on persons without any evidence of pathology is consistent with previous studies in healthy older adults [9,55–57] and in a wide variety of studies regarding populations with various pathologies [29, 30]. Consequently, differences in the brain response pattern between groups with different cognitive reserve levels in both tasks could be interpreted as a function of brain response efficiency [45]. Thus, we expected that the brain electrical response pattern of young people with high CR would reflect a processing that is less vulnerable to the complexity of the task than that of young people with low CR, previously proposed as neural efficiency [9]. Regardless of the type of processing (fluid or crystallized), we think that this positive effect of CR would be observed in the performance of young people for both tasks. There is evidence from neuroimaging studies of greater neural efficiency in cognitively normal adults, such that people with high CR show less task-related activation as a function of increasing task load than people with lower CR [62].

Our results for both experiments showed a modulation of ERP amplitude by WM load on the performance of the young participants that was dependent on the CR level. Similar to what was found in ERP studies using Sternberg-type tasks [6, 9], the low CR group, which was associated with a less efficient performance in both tasks, tended to show a wider distribution of amplitude changes modulated by WM load than those from the high CR group. The P300 latency modulation pattern found supported both our findings and those from previous studies [9]. In contrast, the results of the grammatical judgment task showed that participants with high CR had greater amplitude modulation as a function of WM load. This apparent contrasting result could be interpreted following the same logic regarding efficiency. A low CR does not modulate the LAN and P600a amplitudes as the WM load increases because both WM loads were equally demanding, generating an equally high cost of processing. That is, the low WM load was as costly as the high WM load for the brain responses of subjects with low CR. Our study could suggest that young participants with higher CR have better performance and show greater modulation in the brain response or greater cerebral flexibility. In this way and in accordance with the CR hypothesis [17, 63], the results of this study could suggest that high CR results in the implementation of more-efficient neural networks for the resolution of tasks regardless of the type of processing involved in the execution of the task (fluid or crystallized intelligence).

Research on cognitive reserve has mainly focused on the evaluation of older adults and people with cognitive impairment [55,64,65]. The results of this study support the fact that CR is a malleable entity whose level at any stage of life is dependent on the sum of the experiences. This accumulation of experiences gives the possibility of increasing CR and consequently improving cognitive performance.

Two questions remain to be investigated regarding cognitive reserve. First, which cognitive reserve factors are most important, and which of their interactions contribute to the dynamic processes underlying cognitive/brain development throughout life? What are the more important proxy measures for cognitive reserve, and what can be learned about their beneficial effects?. One of the most-used proxy measures for evaluating CR has been schooling or years of education. In a systematic review of cognitive reserve and its effects on pathological populations, it was shown that in most studies with positive results, the main variable that contributes to building the cognitive reserve is educational level [30]. It should be noted that the years of education for older adults seems to be crystallized, and it maintains its effect throughout their lives. Although older adults have completed their formal studies many years ago, the cognitive reserve has a protective effect. The late-life relations between education and cognitive performance reflect the persistence of education-cognition relations that have existed since earlier adulthood [58]. Thus, our results could show the effects of cognitive reserve in early adulthood.

It has been demonstrated that young and older adults differ in terms of the relationship between brain activity and years of education during a memory task [66]. Increased frontal activity is observed in the less educated young and has been correlated with poor recognition memory. The more educated young adults and those with better memory performance engaged posterior brain regions. Meanwhile, older adults seem to have separate networks related to education and memory performance, with frontal and lateral temporal regions related to more education but not to performance.

## Conclusion

This work provides evidence that cognitive reserve has an effect on other stages of human development in addition to older age. Cognitive reserve can play a modulating role in the performance of cognitive tasks of both fluid and crystalized intelligence processes.

## References

1. Satz P. Brain Reserve Capacity on Symptom Onset After Brain Injury: A Formulation and Review of Evidence for Threshold Theory. Neuropsychology. 1993;7(3):273–95. doi: 10.1037/0894-4105.7.3.273.

2. Stern Y. What is cognitive reserve? Theory and research application of the reserve concept. J Int Neuropsychol Soc. 2002;8(3):448–60. doi: 10.1017.S1355617701020240

3. Pettigrew C, Soldan A. Defining Cognitive Reserve and Implications for Cognitive Aging. Curr Neurol Neurosci Rep. 2019;19(1):1–12. doi: 10.1007/s11910-019-0917-z.

4. Stern Y, Arenaza-Urquijo EM, Bartrés-Faz D, Belleville S, Cantilon M, Chetelat G, et al. Whitepaper: Defining and investigating cognitive reserve, brain reserve, and brain maintenance. Alzheimer’s Dement. 2018;(September):1–7. doi: 10.1016/j.jalz.2018.07.219.

5. Gajewski PD, Wild-Wall N, Schapkin SA, Erdmann U, Freude G, Falkenstein M. Effects of aging and job demands on cognitive flexibility assessed by task switching. Biol Psychol. 2010;85(2):187–99. doi: 10.1016/j.biopsycho.2010.06.009.

6. Gu L, Chen J, Gao L, Shu H, Wang Z, Liu D, et al. Cognitive reserve modulates attention processes in healthy elderly and amnestic mild cognitive impairment: An event-related potential study. Clin Neurophysiol. 2018;129(1):198–207. doi: 10.1016/j.clinph.2017.10.030.

7. Ruiz-Contreras AE, Soria-Rodríguez G, Almeida-Rosas GA, García-Vaca PA, Delgado-Herrera M, Méndez-Díaz M, et al. Low diversity and low frequency of participation in leisure activities compromise working memory efficiency in young adults. Acta Psychol (Amst). 2012 Jan;139(1):91–6. doi: 10.1016/j.actpsy.2011.10.011.

8. Scarmeas N, Levy G, Tang MX, Manly J, Stern Y. Influence of leisure activity on the incidence of Alzheimer’s Disease. Neurology. 2001;57(12):2236–42. doi: 10.1212/wnl.57.12.2236.

9. Speer ME, Soldan A. Cognitive reserve modulates ERPs associated with verbal working memory in healthy younger and older adults. Neurobiol Aging. 2015 Mar;36(3):1424–34. doi: 10.1016/j.neurobiolaging.2014.12.025.

10. Wilson RS, Barnes LL, Bennett DA. Assessment of Lifetime Participation in Cognitively Stimulating Activities. J Clin Exp Neuropsychol. 2003;25(5):634–42. doi: 10.1076/jcen.25.5.634.14572.

11. Angel L, Fay S, Bouazzaoui B, Isingrini M. Individual differences in executive functioning modulate age effects on the ERP correlates of retrieval success. Neuropsychologia. 2010; 48(12): 3540–53. doi: 10.1016/j.neuropsychologia.2010.08.003.

12. Caffò AO, Lopez A, Spano G, Saracino G, Stasolla F, Ciriello G, et al. The role of pre-morbid intelligence and cognitive reserve in predicting cognitive efficiency in a sample of Italian elderly. Aging Clin Exp Res. 2016;28(6):1203–10. doi: 10.1007/s40520-016-0580-z.

13. Fritsch T, McClendon MJ, Smyth K a, Lerner AJ, Friedland RP, Larsen JD. Cognitive functioning in healthy aging: the role of reserve and lifestyle factors early in life. Gerontologist. 2007;47(3):307–22. doi: 10.1093/geront/47.3.307.

14. Le carret N, Lafont S, Mayo W, Fabrigoule C. The effect of education on cognitive performance and its implication for the constitution of the cognitive reserve. Dev Neuropsychol. 2003;23(3):317–337. doi: 10.1207/S15326942DN2303_1.

15. Luerding R, Gebel S, Gebel E-M, Schwab-Malek S, Weissert R. Influence of Formal Education on Cognitive Reserve in Patients with Multiple Sclerosis. Front Neurol. 2016;7(March):1–9. doi: 10.3389/fneur.2016.00046

16. Sumowski JF, Wylie GR, Chiaravalloti N, Deluca J. Intellectual enrichment lessens the effect of brain atrophy on learning and memory in multiple sclerosis. Neurology. 2010;74(24):1942–5. doi: 10.1212/WNL.0b013e3181e396be

17. Stern Y. An approach to studying the neural correlates of reserve. Brain Imaging Behav. 2016;11(2):410–6. doi: 10.1007/s11682-016-9566-x

18. Vasile C. Cognitive Reserve and Cortical Plasticity. Procedia-Soc Behav Sci [Internet]. 2013;78:601–4. doi: 10.1016/j.sbspro.2013.04.359.

19. Zihl J, Fink T, Pargent F, Ziegler M, Bühner M. Cognitive reserve in young and old healthy subjects: differences and similarities in a testing-the-limits paradigm with DSST. PLoS One. 2014 Jan;9(1):e84590. doi: 10.1371/journal.pone.0084590.

20. Cabeza R, Daselaar SM, Dolcos F, Prince E, Budde M, Nyberg L. Task-independent and Task-specific Age Effects on Brain Activity during Working Memory, Visual Attention and Episodic Retrieval. 2004;(April):364–75. doi: 10.1093/cercor/bhg133.

21. Scarmeas N, Zarahn E, Anderson KE, Hilton J, Flynn J, Van Heertum RL, et al. Cognitive reserve modulates functional brain responses during memory tasks: a PET study in healthy young and elderly subjects. Neuroimage. 2003 Jul;19(3):1215–27. doi: 10.1016/s1053-8119(03)00074-0.

22. Stern Y, Habeck C, Moeller J, Scarmeas N, Anderson KE, Hilton HJ, et al. Brain networks associated with cognitive reserve in healthy young and old adults. Cereb Cortex. 2005 Apr;15(4):394–402. doi: 10.1093/cercor/bhh142.

23. Stern Y, Zarahn E, Habeck C, Holtzer R, Rakitin BC, Kumar A, et al. A common neural network for cognitive reserve in verbal and object working memory in young but not old. Cereb Cortex. 2008 Apr;18(4):959–67. doi: 10.1093/cercor/bhm134.

24. Chillemi G, Scalera C, Terranova C, Calamuneri A, Buccafusca M, Dattola V, et al. Cognitive processess and cognitive reserve in multiple sclerosis. Arch Ital Biol. 2015;153:19–24. doi: 10.4449/aib.v153i1.3696.

25. Sandry J, Sumowski JF. Working Memory Mediates the Relationship between Intellectual Enrichment and Long-Term Memory in Multiple Sclerosis: An Exploratory Analysis of Cognitive Reserve. J Int Neuropsychol Soc. 2014 Jul 14;20(8):1–5. doi: 10.1017/S1355617714000630.

26. Cabeza R. Hemispheric Asymmetry Reduction in Older Adults : The HAROLD Model. Psychol Aging. 2002;17(1):85–100. doi: 10.1037//0882-7974.17.1.85.

27. Davis SW, Dennis NA, Daselaar SM, Fleck MS, Cabeza R. Qué PASA? the posterior-anterior shift in aging. Cereb Cortex. 2008;18(5):1201–9. doi: 10.1093/cercor/bhm155.

28. Park DC, Reuter-Lorenz P. The Adaptive Brain: Aging and Neurocognitive Scaffolding. Annu Rev Psychol. 2009;60:173–96. doi: 10.1146/annurev.psych.59.103006.093656.

29. Baxendale S, Heaney D, Rugg-Gunn F, Friedland D. Neuropsychological outcomes following traumatic brain injury. Pract Neurol. 2019;0:1–7. doi: 10.1136/practneurol-2018-002113.

30. Reynoso-Alcántara V, Silva-Pereyra J, Fernández-Harmony T, Mondragón-Maya A. Principales efectos de la reserva cognitiva sobre diversas enfermedades: una revisión sistemática. Psiquiatr Biológica. 2018;(October). doi: 10.1016/j.psiq.2018.02.005

31. Sundgren M, Wahlin Å, Maurex L, Brismar T. Event related potential and response time give evidence for a physiological reserve in cognitive functioning in relapsing–remitting multiple sclerosis. J Neurol Sci. 2015;356(1–2):107–12. doi: 10.1016/j.jns.2015.06.025

32. Donchin E. Surprise!…Surprise? Psychophysiology. 1981;18(5): 493–513. doi: 10.1111/j.1469-8986.1981.tb01815.x.

33. Polich J. Updating P300: An integrative theory of P3a and P3b. Clin Neurophysiol. 2007;118(10):2128–48. doi: 10.1016/j.clinph.2007.04.019.

34. Kok A. On the utility of P300 amplitude as a measure of processing capacity. Psychophysiology. 2001;38:557–577. doi: 10.1017/S0048577201990559

35. Polich J, Corey-Bloom J. Alzheimers Disease and P300: Review and Evaluation of Task and Modality. Curr Alzheimer Res. 2005;2(5):515–25. doi: 10.2174/156720505774932214

36. Chang Y-K, Huang C-J, Chen K-F, Hung T-M. Physical activity and working memory in healthy older adults: An ERP study. Psychophysiology. 2013;50(11):1174–82. doi: 10.1111/psyp.12089

37. Buchmann J, Gierow W, Reis O, Haessler F. Intelligence moderates impulsivity and attention in ADHD children: An ERP study using a go/nogo paradigm. World J Biol Psychiatry. 2011;12(SUPPL. 1):35–9. doi: 10.3109/15622975.2011.600354.

38. Wronka E, Kaiser J, Coenen AML. Psychometric intelligence and P3 of the event-related potentials studied with a 3-stimulus auditory oddball task. Neurosci Lett. 2013;535(1):110–5. doi: 10.1016/j.neulet.2012.12.012

39. Stine-Morrow EA, Soederberg-Miller LM, Gagne DD, Hertzog C. Self-Regulated Reading in Adulthood. Psychol Aging. 2008;23(1):131–53. doi: 10.1037/0882-7974.23.1.131.

40. Payne BR, Gao X, Noh SR, Anderson CJ, Stine-Morrow EA. The effects of print exposure on sentence processing and memory in older adults: Evidence for efficiency and reserve. Neuropsychol Dev Cogn B Aging Neuropsychol Cogn. 2012;19(1–2):122–49. doi: 10.1080/13825585.2011.628376.

41. Alatorre-Cruz GC, Silva-Pereyra J, Fernández T, Rodríguez-Camacho MA, Castro-Chavira SA, Sanchez-López J. Effects of Age and Working Memory Load on Syntactic Processing : An Event-Related Potential Study. Front Hum Neurosci. 2018;12(May):1–11. doi: 10.3389/fnhum.2018.00185.

42. Friederici AD. Towards a neural basis of auditory sentence processing. Trendd Cogn Sci. 2002;6(2):78–84. doi: 10.1016/s1364-6613(00)01839-8.

43. Molinaro N, Barber HA, Carreiras M. Grammatical agreement processing in reading: ERP findings and future directions. Cortex. 2011;47(8):908–30. doi: 10.1016/j.cortex.2011.02.019

44. Barber H, Carreiras M. Grammatical Gender and Number Agreement in Spanish: An ERP Comparison. J Cogn Neurosci. 2005;17(1):137–53. doi: 10.1162/0898929052880101

45. Barulli D, Stern Y. Efficiency, capacity, compensation, maintenance, plasticity: emerging concepts in cognitive reserve. Trends Cogn Sci. 2013;17(10):502–9. doi: 10.1016/j.tics.2013.08.012.

46. Farage MA, Osborn TW, MacLean AB. Cognitive, sensory, and emotional changes associated with the menstrual cycle: A review. Arch Gynecol Obstet. 2008;278(4):299–307. doi: 10.1007/s00404-008-0708-2.

47. Instituto Nacional de Estadística y Geografía. INEGI. Sistema nacional de clasificación de ocupaciones SINCO. 2011. 180 p.

48. Wechsler D. Escala de Inteligencia para Adultos-IV (WAIS-IV). México: El Manual Moderno; 2014.

49. Dotson VM, Schinka JA, Brown LM, Mortimer JA, Borenstein AR. Characteristics of the Florida Cognitive Activities Scale in Older African Americans. Assessment. 2009;15(1):72–7. doi: 10.1177/1073191107307509.

50. Nucci M, Mapelli D, Mondini S. Cognitive Reserve Index questionnaire (CRIq): a new instrument for measuring cognitive reserve. Aging Clin Exp Res. 2012;24(3):218–26. doi: 10.3275/7800

51. Valenzuela M, Sachdev P. Assessment of complex mental activity across the lifespan : development of the Lifetime of Experiences Questionnaire (LEQ). Psychol Med. 2007;37(7):1015–25. doi: 10.1017/S003329170600938X.

52. Luck SJ. An Introduction to the Event-Related Potentials Technique. Second edit. United States: MIT Press; 2014.

53. Pascual-Marqui RD, Lehmann D, Koukkou M, Kochi K, Anderer P, Saletu B, et al. Assessing interactions in the brain with exact low-resolution electromagnetic tomography Assessing interactions in the brain with exact low-resolution electromagnetic tomography. Philos Trans A Math Phys Eng Sci. 2011;369(1952):3768–84. doi: 10.1098/rsta.2011.0081.

54. Benjamini Y, Hochberg Y. Benjamini Y, Hochberg Y. Controlling the false discovery rate: a practical and powerful approach to multiple testing. J R Stat Soc B. 1995;57(1):289–300. doi: 10.2307/2346101.

55. Opdebeeck C, Martyr A, Clare L. Cognitive reserve and cognitive function in healthy older people: A meta-analysis. Neuropsychol Dev Cogn B Aging Neuropsychol Cogn 2016;23(1):40–60. doi: 10.1080/13825585.2015.1041450.

56. Roldán-Tapia L, García J, Cánovas R, León I. Cognitive reserve, age, and their relation to attentional and executive functions. Appl Neuropsychol Adult. 2012 Jan;19(1):2–8. doi: 10.1080/09084282.2011.595458.

57. Soto-Añari M, Flores-Valdivia G, Fernández-Guinea S. Nivel de lectura como medida de reserva cognitiva en adultos mayores. Rev Neurol. 2013;2(56):79–85. doi: 10.33588/rn.5602.2012402.

58. Tucker-Drob EM, Johnson KE, Jones RN. The Cognitive Reserve Hypothesis: A Longitudinal Examination of Age-Associated Declines in Reasoning and Processing Speed. Dev Psychol. 2009;45(2):431–46. doi:10.1037/a0014012.

59. Friederici AD. The time course of syntactic activation during language processing: A model based on neuropsychological and neurophysiological data. Brain Lan. 1995; 50(3):259–81 doi: 10.1006/brln.1995.1048.

60. Kolk HHJ, Chwilla DJ, Herten M Van, Oor PJW. Structure and limited capacity in verbal working memory : A study with event-related potentials. 2003;85(1):1–36 doi: 10.1016/s0093-934x(02)00548-5.

61. Kluender R, Kutas M. Bridging the Gap : Evidence & om ERPs on the Processing of Unbounded Dependencies. J Cogn Neurosci. 1993;5(2):196–214. doi: 10.1162/jocn.1993.5.2.196.

62. Habeck C, Hilton HJ, Zarahn E, Flynn J, Moeller J, Stern Y. Relation of cognitive reserve and task performance to expression of regional covariance networks in an event-related fMRI study of nonverbal memory. Neuroimage. 2003 Nov;20(3):1723–33. doi: 10.1016/j.neuroimage.2003.07.032.

63. Stern Y. Cognitive reserve. Neuropsychologia. 2009;47(10):2015–28. doi: 10.1016/j.neuropsychologia.2009.03.004.

64. Hindle JV., Martyr A, Clare L. Cognitive reserve in Parkinson’s disease: A systematic review and meta-analysis. Parkinsonism Relat Disord. 2014;20(1):1–7. doi: 10.1016/j.parkreldis.2013.08.010.

65. Stern Y, Gurland B, Tatemichi TK, Tang MX, Wilder D, Mayeux R. Influence of education and occupation on the incidence of Alzheimer’s disease. JAMA. 1994;271(13):1004–10. doi: 10.1001/jama.1994.03510370056032.

66. Springer MV, McIntosh AR, Winocur G, Grady CL. The relation between brain activity during memory tasks and years of education in young and older adults. Neuropsychology. 2005;19(2):181–92. doi: 10.1037/0894-4105.19.2.181.

